# Copy number motifs expose genome instability type and predict driver events and disease outcome in breast cancer

**DOI:** 10.1101/769356

**Authors:** Arne V. Pladsen, Gro Nilsen, Oscar M. Rueda, Miriam R. Aure, Ørnulf Borgan, Knut Liestøl, Valeria Vitelli, Arnoldo Frigessi, Anita Langerød, OSBREAC, Anthony Mathelier, Olav Engebråten, David C. Wedge, Peter Van Loo, Carlos Caldas, Anne-Lise Børresen-Dale, Hege G. Russnes, Ole Christian Lingjærde

**Affiliations:** Department of Cancer Genetics, Institute for Cancer Research, Oslo University Hospital, Oslo, Norway; Centre of Bioinformatics, Department of Informatics, University of Oslo, Oslo, Norway; Cancer Research UK, Cambridge Research Institute, Cambridge, UK; Department of Mathematics, University of Oslo, Oslo, Norway; Institute of Basic Medical Sciences, Faculty of Medicine, University of Oslo, Oslo, Norway; The Oslo Breast Cancer Consortium; Centre for Molecular Medicine Norway, University of Oslo, Oslo, Norway; Big Data Institute, Li Ka Shing Centre for Health Information and Discovery, University of Oxford, UK; NIHR Biomedical Research Centre, Oxford, UK; The Francis Crick Institute, London, UK; Department of Human Genetics, University of Leuven, Leuven, Belgium; Institute for Clinical Medicine, University of Oslo, Oslo, Norway; Department of Pathology, Oslo University Hospital, Oslo, Norway; KG Jebsen Centre for B-cell malignancies, Institute for Clinical Medicine, University of Oslo, Norway

## Abstract

Tumor evolution is dependent on and constrained by the genotypes emerging from genome instability. We hypothesized that non-site-specific copy number motifs would correlate with underlying replication defects and also with tumor and patient fate. Six feature detectors were defined to characterize and score the local spatial behaviour of a copy number profile. By accumulating scores across genomic regions, a low-dimensional representation of the tumor genome was obtained. The proposed Copy Aberration Regional Mapping Analysis (CARMA) algorithm was applied to 2384 breast tumors from three breast cancer cohorts, revealing distinct copy number motifs in established molecular subtypes. A prognostic index combining the features predicted breast cancer specific survival better than both the genomic instability index (GII) and all commonly used clinical stratifications. CARMA offers effective comparison of tumor subgroups and extracts biologically and clinically relevant features from allele-specific copy number profiles.

## 1 Introduction

The allele-specific copy number profile of a tumor is a window into its past history and its future evolutionary potential [1, 2, 3]. A range of different mechanisms and defects may be involved, including aneuploidy, genomic duplication, deletion, inversion, translocation, double minute chromosomes, breakage-fusion-bridge cycles, chromothripsis and chromoplexy [4, 5, 6, 7]. Depending on the events and their cause, distinct traces may be left on the copy number level, including the presence of global aberration patterns such as simplex, complex and sawtooth [8, 9].

Several algorithms have been proposed to detect locus specific or type specific copy number aberrations in tumors. The chromosomal instability index (CINdex) [10] and the genomic instability index (GII) [11] both quantify the total amount of genomic aberrations. Stratifications aiming to identify specific aberration patterns are also available, including locus specific aberrations (GISTIC) [12, 13], simplex and complex copy number events [9] and structural rearrangement patterns [14].

In general, we may consider a copy number profile as consisting of both site-specific events and more general regional features (motifs) present throughout the genome. Here, we hypothesize that such motifs represent a substantial proportion of the copy number variation in a tumor and also partly explains the high inter-tumor copy number heterogeneity frequently observed in cancer. We further hypothesize that the presence or absence of specific motifs is informative of a tumor’s past and future evolutionary trajectory. Detailed characterization of such features would thus allow prediction of disease behaviour and could potentially direct choice of treatment. The potential link between such features and DNA repair defects and mutational processes has recently been demonstrated in ovarian cancer [15].

Here, we present a computational framework for extraction and analysis of regional non-site specific motifs from allele-specific copy number profiles. The core of this framework is the Copy Aberration Regional Mapping Analysis (CARMA) algorithm, which creates a compact representation of the aberration architecture. Conceptually, the algorithm represents copy number profiles as real-valued functions over the genomic domain and derives a small set of scores representing distinct regional features. CARMA can be viewed as an extension of the chromosomal and genomic instability indices that also takes into account copy number amplitude, distribution of copy number break points (including spatial distribution along the chromosome) and allelic imbalance. It captures the degree of regional fluctuations in copy number, a signature feature of e.g. chromothripsis and chromoplexy. By generating a low-dimensional representation of the copy number data, the proposed algorithm also avoids the ‘curse of dimensionality’.

To demonstrate that the proposed method captures biologically and clinically relevant characteristics, the method was applied to three different breast cancer data sets (METABRIC, Oslo2, and OsloVal). A novel prognostic index integrating all six modes of genomic aberrations is derived and is shown to predict breast cancer specific survival better than commonly used clinical variable stratifications. The proposed index also outperforms the genomic instability index GII. We demonstrate the relation between the copy number motifs and established molecular and clinical expression markers and the relation to driver gene based classifications of breast cancer.

CARMA is applicable to allele-specific copy number data from any type of platform, including SNP arrays and high-throughput sequencing. Software will be made available on GitHub.

## 2 Results

### 2.1 Brief outline of the analysis approach

CARMA is applicable to allele-specific copy number profiles from one or several tumors, obtained from SNP array analysis or DNA high-throughput sequencing. The analysis pipeline is depicted in Figure 1a. First, the algorithm extracts multiple local features, and next these are accumulated across genomic regions by integration to form six numerical regional scores (Figure 1b). Precise mathematical definitions are deferred to Material and Methods. The extracted features detect generic variational properties of functions and link closely to chromothripsis, loss of heterozygosity, and genomic loss and gain (Figure 1c). An application of the algorithm to three breast tumor samples in the Oslo2 cohort and with chromosome arms as regions is shown in Figure 1d. Specific regional features are discernible, illustrating how CARMA can be used to perform between-sample comparison of copy number features that are not locus specific.

**Figure 1.**
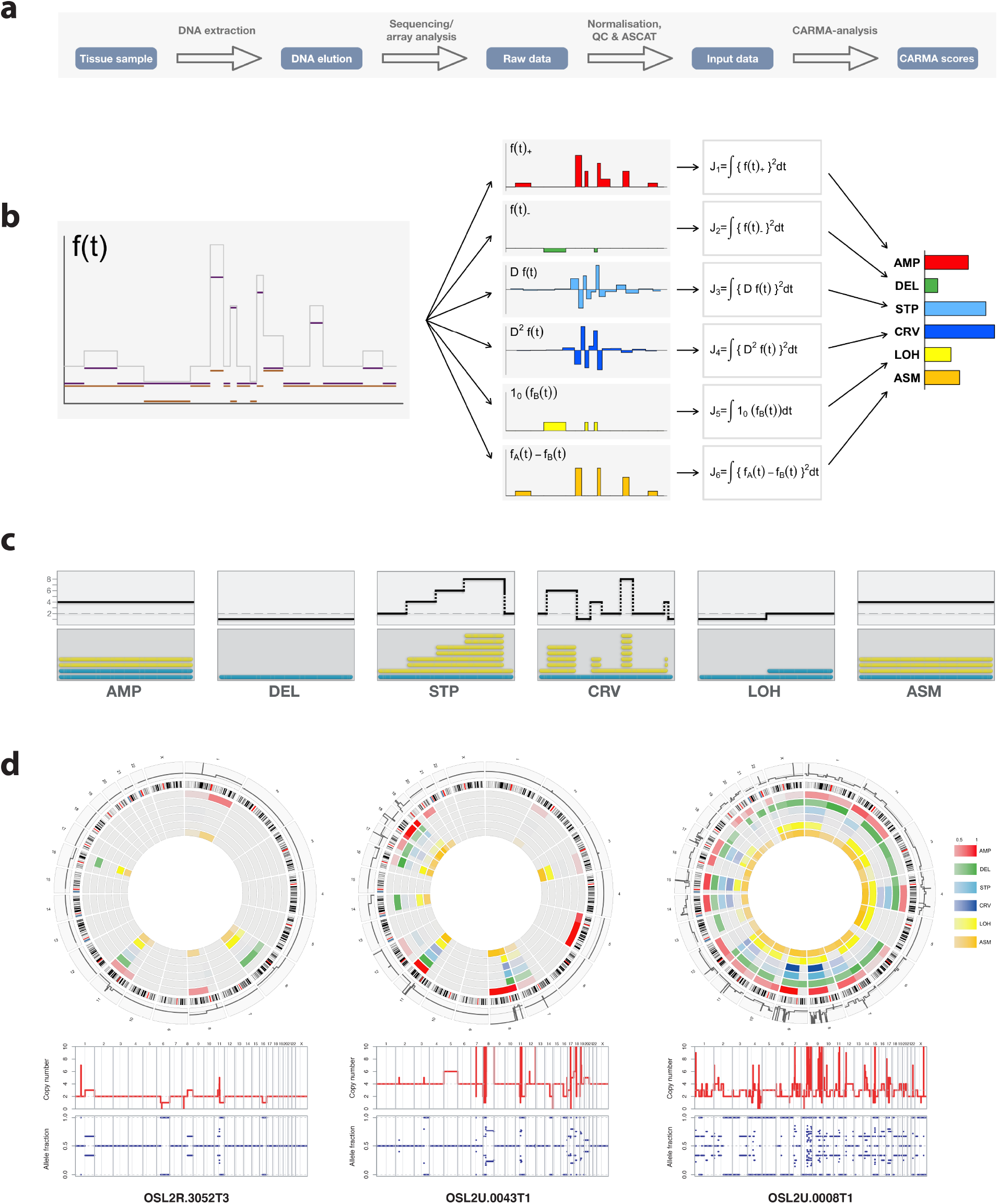
Outline of the CARMA algorithm. **a** Analysis pipeline. **b** Construction of regional CARMA scores. The input is one or more allele-specific copy number profiles. The algorithm extracts local features and accumulates these across genomic regions to form six regional scores. **c** Prototype patterns captured by each of the six CARMA scores. **d** An application of the algorithm to three breast tumor samples in the Oslo2 cohort. Lower panel: total copy number (logR values) and allele fraction as a function of genomic locus. Upper panel: Circos plots of regional (arm-wise) CARMA scores.

### 2.2 Molecular subgroups have distinct CARMA signatures

We next considered the distribution of CARMA scores within established molecular stratifications of breast carcinomas (PAM50 and IntClust). PAM50 [16, 17] is an expression based classification system defining five distinct subgroups of breast tumours based on the correlation to a set of 50 genes. IntClust [1, 18] identifies ten different subtypes based on the pattern of copy number aberrations exerting an effect on gene expression *in cis*. The distribution of CARMA scores within these classification systems were explored in three different breast cancer data sets of varying sample size (*n* = 1943, *n* = 276, and *n* = 165). The percentage of tumors with scores exceeding a median threshold was plotted for all arm scores and for each PAM50 and IntClust subtype separately (Figure 2a and Supplementary Figure 1-3). The CARMA scores consistently reflected differences in the landscapes of genomic architecture in the different biological and clinical patient groups. This visual overview of aberration patterns highlights subtype specific features such as frequent allelic loss on 17p and frequent gain and high complexity on 17q in IntClust1; gain on 1q, frequent asymmetric gain and complex aberrations on 11q and allelic loss on 16q in IntClust2; etc. The signatures of regional CARMA scores within the PAM50 subtypes highlight known features, including whole arm 1q gain/16q loss in luminal A tumours, the more complex copy number aberrations in luminal B tumours, the 17q alterations dominating Her2-enriched tumours and the global instability of basal-like tumours. Three-dimensional scatter plots of CARMA scores were plotted for all tumors in the Oslo2 cohort (*n* = 276) and METABRIC cohort (*n* = 1943) (see Figure 2b). Trend curves and subtype centroids both demonstrate high degree of consistency between the two cohorts.

**Figure 2.**
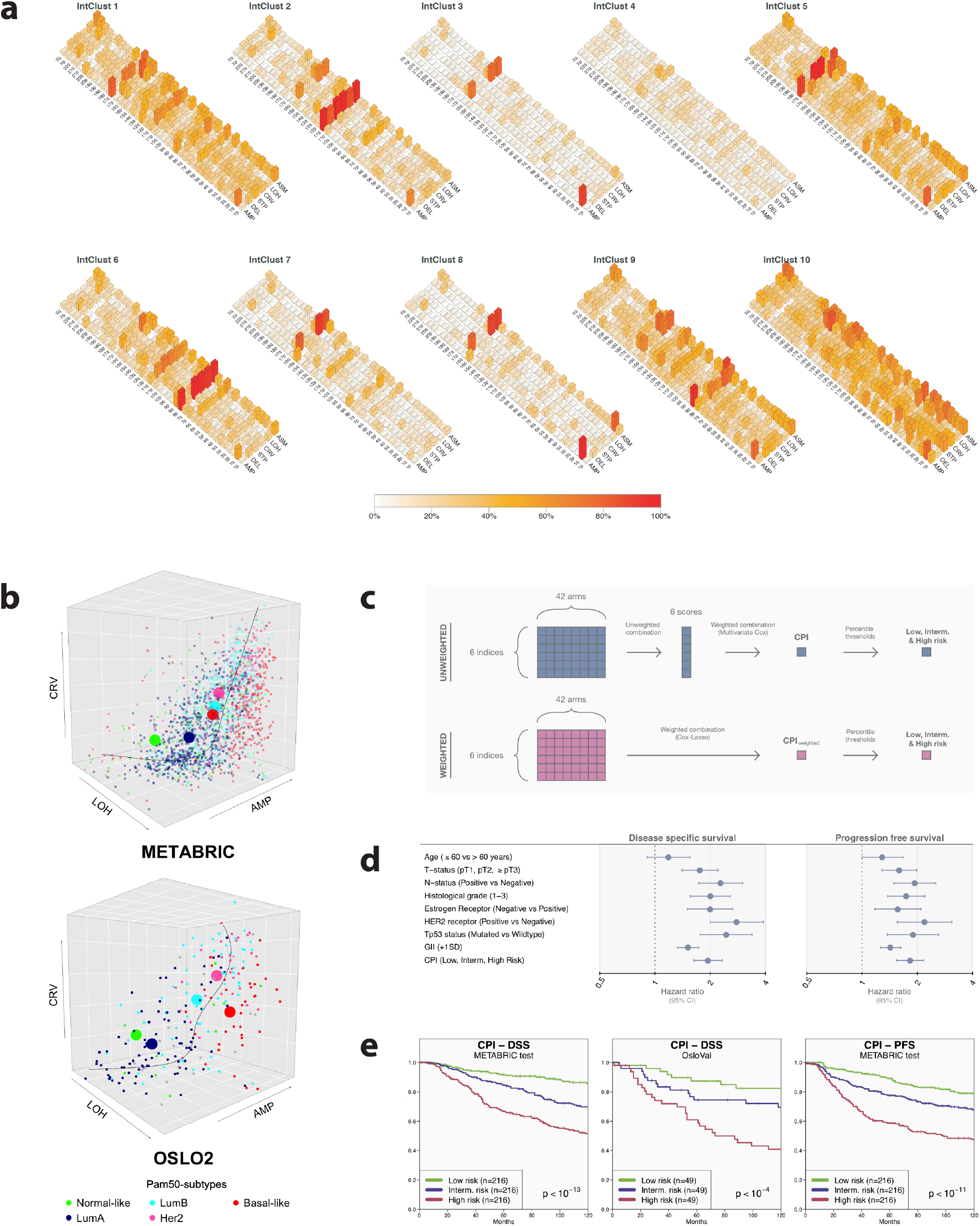
Stratification and outcome prediction with CARMA. **a** CARMA score distribution within each of the 10 IntClust subtypes defined in Curtis et al.[1]. The height of each bar represents the proportion of samples in the subgroup with an arm score above the median score for this index across all arms and subtypes. **b** Three-dimensional scatter plots of tumors using three of the CARMA scores designed to detect three major categories of copy number aberration patterns in tumors (amplifications: AMP, allelic loss: LOH and complex rearrangements: CRV). Colors indicate PAM50 subtype (see legend at bottom) and large spheres show subtype centroids. Upper panel: Oslo2 cohort (n = 276); Lower panel: METABRIC cohort (n = 1943). **c** Flow chart depicting the construction of prognostic indices (CPIs) from the arm-wise CARMA scores, based on the METABRIC discovery cohort. Upper panel: Construction of the CPI. Arm-wise scores are first collapsed by taking an unweighted average, and the resulting six genome-wide scores are combined by multivariate Cox regression. Thresholds corresponding to the 1/3 and 2/3 percentile were applied to classify samples into groups of low, intermediate and high risk, with numerical values ranging from 1-3. Lower panel: Construction of the CPI_*weighted*_. Arm-wise scores are combined by cross-validated multivariate Cox-Lasso regression, resulting in one genome-wide score. Thresholds corresponding to the 1/3 and 2/3 percentile were applied as above to classify samples into groups of low, intermediate and high risk. **d** Hazard ratios and 95% confidence intervals(CI) for clinical variables, the CARMA prognostic index(CPI) based on unweighted averages of arm-wise CARMA scores, and the genomic instability index(GII). Shown are unadjusted estimates for disease specific survival (DSS) and progression free survival (PFS). **e** Survival prediction using the CPI stratified into low, intermediate and high risk groups. Kaplan-Meier plots of disease specific survival (DSS) for the three risk stratifications within the METABRIC test and OsloVal set, as well as for progression free survival within the METABRIC test set. From left to right: DSS METABRIC test cohort; DSS OsloVal cohort; PFS METABRIC test cohort.

### 2.3 Predicting survival from regional scores

To assess the association between breast cancer specific survival and genome-wide CARMA scores, a univariate Cox proportional hazards regression model was fitted with each score as a covariate (see Supplementary Table 3). For this purpose we used the largest cohort (METABRIC set). All scores were associated with survival (*P <* 10^*−*6^) and the strongest associations were found for the genes *STP* and *CRV* (*P <* 10^*−*15^).

We next split the METABRIC cohort into a discovery cohort (*n* = 1295) and a test cohort (*n* = 648). We fitted a multivariate Cox regression model to disease specific survival (DSS) and progression free survival (PFS) data in the discovery cohort based on the six predictors. The predictors were defined by taking an unweighted mean across all the regional (arm-wise) CARMA scores (Figure 2c). The fitted model was next applied to the test set, producing a single unweighted prognostic value per patient. Thresholds corresponding to the 1/3 and 2/3 percentile were applied to classify samples into groups of low, intermediate and high risk, with numerical values ranging from 1-3. This final score was termed the CARMA Prognostic Index (CPI). An alternative prognostic index was defined using the 252 armwise CARMA scores directly as predictors and fitting a Cox regression model with Lasso penalty to the training set. Coefficients derived from the analysis (Supplementary Figure 4) were used as weights to calculate a weighted prognostic index termed CPI_*weighted*_.

To compare the efficacy of the CPI and CPI_*weighted*_ to established clinically and biologically relevant parameters, we fitted a univariate Cox regression model in the METABRIC test set using the prognostic indices and the clinical parameters as covariates (Table 1 and Supplementary Table 4-5). The P-value for the CPI from the analysis was lower than for any of the other clinical parameters when looking at both DSS and PFS (*P* = 2.4 ⋅ 10^*−*13^ and *P* = 3.8 ⋅ 10^*−*12^ for DSS and PFS respectively), and also performed better than the CPI_*weighted*_. The CPI_*weighted*_ did however remain strongly significant in the analysis (*P* = 2.2 ⋅ 10^*−*7^ and *P* = 6.6 ⋅ 10^*−*9^ for DSS and PFS respectively) presenting P-values lower than many of the other established parameters.

**Table 1.** Prognostic value of clinical variables, the CARMA prognostic index(CPI), and the genomic instability index(GII), for disease specific survival (DSS) and progression free survival (PFS). HR: hazard ratio.

| Type | Covariate | Disease specific survival |  | Progression free survival |  |
| --- | --- | --- | --- | --- | --- |
|  |  | P value | HR | P value | HR |
| Unadjusted | Age ($\leq 60$ vs $> 60$ years) | 2.2e-01 | 1.19 | 5.3e-02 | 1.29 |
| Unadjusted | T-status (pT1, pT2, ≥pT3) | 9.2e-07 | 1.76 | 2.9e-05 | 1.59 |
| Unadjusted | N-status (Positive vs Negative) | 5.5e-09 | 2.28 | 8.2e-07 | 1.94 |
| Unadjusted | Histological grade (1-3) | 5.9e-08 | 2.01 | 3.2e-06 | 1.74 |
| Unadjusted | Estrogen Receptor (Negative vs Positive) | 2.0e-06 | 2.00 | 2.3e-03 | 1.57 |
| Unadjusted | HER2 receptor (Positive vs Negative) | 1.6e-09 | 2.79 | 7.0e-06 | 2.19 |
| Unadjusted | Tp53 status (Mutated vs Wildtype) | 7.2e-08 | 2.44 | 1.0e-04 | 1.90 |
| Unadjusted | GII ($+1SD$) | 1.2e-09 | 1.51 | 3.9e-08 | 1.43 |
| Unadjusted | CPI (Low,Interm.,High risk) | 2.4e-13 | 1.95 | 3.8e-12 | 1.83 |
| Adjusted | CPI (Low,Interm.,High risk) | 4.5e-13 | 1.94 | 1.1e-11 | 1.81 |
| | Age ($\leq 60$ vs $> 60$ years) | 5.8e-01 | 1.08 | 2.1e-01 | 1.18 |
| Adjusted | CPI (Low,Interm.,High risk) | 6.8e-12 | 1.88 | 6.2e-11 | 1.78 |
| | T-status (pT1, pT2, ≥pT3) | 1.0e-05 | 1.68 | 4.7e-04 | 1.49 |
| Adjusted | CPI (Low,Interm.,High risk) | 7.3e-13 | 1.93 | 8.7e-12 | 1.82 |
|  | N-status (Positive vs Negative) | 1.8e-08 | 2.22 | 2.0e-06 | 1.90 |
| Adjusted | CPI (Low,Interm.,High risk) | 2.0e-08 | 1.74 | 2.5e-08 | 1.71 |
|  | Histological grade (1-3) | 1.8e-03 | 1.53 | 2.8e-02 | 1.32 |
| Adjusted | CPI (Low,Interm.,High risk) | 2.9e-11 | 1.86 | 8.7e-11 | 1.79 |
|  | Estrogen Receptor (Negative vs Positive) | 1.4e-03 | 1.61 | 1.7e-01 | 1.23 |
| Adjusted | CPI (Low,Interm.,High risk) | 8.1e-10 | 1.79 | 1.2e-09 | 1.73 |
|  | HER2 receptor (Positive vs Negative) | 4.1e-04 | 1.88 | 3.2e-02 | 1.48 |
| Adjusted | CPI (Low,Interm.,High risk) | 3.7e-05 | 1.64 | 4.2e-05 | 1.60 |
|  | Tp53 status (Mutated vs Wildtype) | 3.4e-03 | 1.71 | 1.2e-01 | 1.33 |
| Adjusted | CPI (Low,Interm.,High risk) | 1.4e-06 | 1.75 | 3.2e-06 | 1.72 |
| | GII ($+1SD$) | 1.2e-01 | 1.15 | 4.2e-01 | 1.07 |

Cox regression models with the prognostic indices as predictors and adjusting for available clinical variables were also considered (Table 1 and Supplementary Table 4-5). The CPI consistently showed smaller P-values than all other clinical variables. The CPI_*weighted*_ also remained significant when adjusting for other variables (Supplementary Table 4-5). Corresponding Hazard ratios for CPI is shown in Figure 2d and Supplementary Figure 5.

The CPI was used to stratify patients into low, intermediate and high-risk groups in the two validation cohorts with survival data available (METABRIC test set and OsloVal). A logrank test was performed for the three groups in each data set (Figure 2e). P-values were significant when considering both DSS (*P <* 10^*−*13^ in METABRIC test and *P <* 10^*−*4^ in OsloVal) and PFS (*P <* 10^*−*11^ in METABRIC test; PFS data was not available for OsloVal).

Finally, the unweighted continuous prognonstic score that was used to obtain the CPI, was utilized to calculate a Harrell’s C score inn the METABRIC test set. The C scores obtained from the analysis were 0.65 and 0.64 based on DSS and PFS respectively.

## 3 Methods and Materials

### Deriving allele-specific copy number profiles

Affymetrix CEL-files were preprocessed using the PennCNV libraries for Affymetrix data [19] that includes quantile normalization, signal extraction and summarization. All samples were normalized to a collection of around 5000 normal samples from the HapMap project [20], the 1000 genome project [21] and the Wellcome Trust Case Control Consortium [22]. The resulting LogR and BAF (B allele frequency) values were segmented with the PCF (piecewise constant fitting) algorithm [23] and processed with the ASCAT algorithm (version 2.3) [24] after adjusting LogR for GC binding artifacts [25]. ASCAT infers an allele specific copy number profile of a tumor after correction for tumor ploidy and tumor cell fraction, and is based on allele specific segmentation of normalized raw data [23] with penalty parameter (*γ*) set to 50. The profile reflects the copy number state at *m* genomic loci for which two alleles are present in the germline in the general population, and can be represented as a sequence of pairs (*n*_*Ai*_, *n*_*Bi*_) (*i* = 1*, …, m*) where *n*_*Ai*_ and *n*_*Bi*_ denote the number of copies of each of two alleles (here called A and B) being present in the tumor genome at the *i*th locus. Pairs are ordered according to location, and since the labels A and B are arbitrary, we may assume that *n*_*Ai*_ ≥ *n*_*Bi*_.

### Calculating regional instability scores

We characterize the allele specific copy number in a small genomic neighborhood on a chromosome arm by six features: degree of alteration in negative direction, degree of alteration in positive direction, degree of change, degree of oscillation, extent of loss of heterozygosity, and extent of allelic imbalance (see Figure 1c). Sliding the genomic region along the chromosome arm from one end to the other, we may regard each feature as a function of genomic position. Specifically, suppose we have measured allele-specific copy numbers (*n*_*Ai*_, *n*_*Bi*_) at genomic loci *L*_*i*_, *i* = 1, …, *m*. We can represent this as a pair of piecewise constant functions (*f*_*A*_, *f*_*B*_) defined on the unit interval *R* = [0, 1]. The interpretation of this is that we have a one-to-one correspondence between *t ∈* [0, 1] and genomic loci *L*(*t*), and if *L*_*k*_ is the measurement locus closest to *L*(*t*), then *f*_*A*_(*t*) = *n*_*Ak*_ and *f*_*B*_(*t*) = *n*_*Bk*_. We assume that *f*_*B*_(*t*) *≤ f*_*A*_(*t*) for all *t ∈ R*, i.e. B is the minor allele when allelic imbalance is present. The median centred total copy number in locus *t* is *f* (*t*) = *f*_*A*_(*t*) + *f*_*B*_(*t*) *− m*, where *m* is the least number in Range(*f*) that satifies *µ*(*f ^−^*^1^((*−∞, m*])) *≥* 1*/*2, where *µ* is the Lebesgue measure. Informally, this means that *m* is chosen as the observed copy number with the property that half the genome has a total copy number less than or equal to *m*. We define the change in total copy number as the derivative *Df* (*t*) of the first order spline interpolation to the center points of segments in *f*, i.e. *Df* (*t*) is the slope of the line segment connecting the pair of segment centers immediately to the left and right of position *t*. Note that *Df* is also a piecewise constant function. We define the oscillation in total copy number as *D*^2^*f* (*t*) = *D*(*Df* (*t*)), which is also a piecewise constant function. This process can in principle be repeated to define higher order properties of *f* such as *D*^3^*f* (*t*) = *D*(*D*^2^*f* (*t*)); however, in practice further levels add little additional information.

Regional instability scores are next defined by integrating the above local scores over the desired region (e.g. over a chromosome arm). To assess the degree of positive or negative deviation within a region, we define two scores:

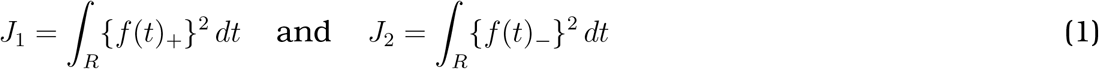

where *z*_+_ = *z* if *z >* 0 and *z*_+_ = 0 otherwise, and *z_−_* = *z* if *z <* 0 and *z_−_* = 0 otherwise. For example, in a region with total copy number equal to the median, we have *J*_1_ = *J*_2_ = 0, while in a region with some gains and no losses relative to the median, we have *J*_1_ *>* 0 and *J*_2_ = 0. The regional degree of change and oscillation in copy number are captured by the following two scores:

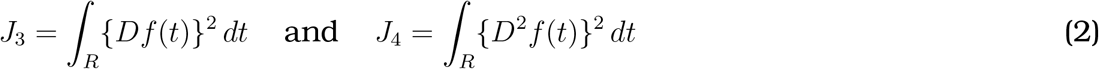

In a region with constant total copy number, we have *J*_3_ = *J*_4_ = 0. In a region with gradually increasing (or decreasing) copy number, *J*_3_ *>* 0 while *J*_4_ is close to zero, and in a region with fluctuations between smaller and larger copy numbers we have *J*_3_ *>* 0 and *J*_4_ *>* 0. Loss of heterozygosity and allelic asymmetry are captured by the last two scores:

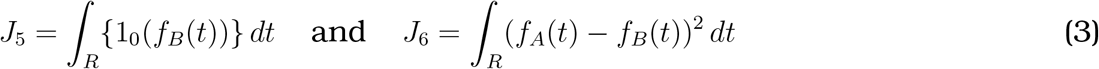

where 1_0_(*z*) = 1 if *z* = 0 and 1_0_(*z*) = 0 otherwise. In a region with only one allele present we have *J*_5_ *>* 0 and the magnitude of the score reflects the proportion of the region with loss of heterozygosity. In a region with allelic imbalance, we have *J*_6_ *>* 0. Further computational details can be found in Supplementary Materials.

### 3.1 Materials

The data material in this study was obtained from three different patient cohorts: METABRIC (n=1943), Oslo2 (n=276) and OsloVal (n=165). The distribution of clinical parameters within each of the datasets can be found in Supplementary Tables 1-2. The METABRIC cohort was randomly split in a 2:1 ratio into a discovery set (n=1295) and a test set (n=648) for the purpose of model validation. See webpage for detailed information regarding sample stratification. For more details about the three cohorts, see Supplementary Material and Methods. Survival data were not available for the Oslo2 cohort. The Regional Comittee for Medical and Health Research Ethics for southeast Norway has approved the study (approval number 1.2006.1607, amendment 1.2007.1125).

## 4 Discussion

Structural DNA distortions are a result of deregulated DNA repair and maintenance, and mutagenic processes operating in the cells. The conventional focus in studies of DNA copy number alterations in tumors is the identification of recurrently deleted and amplified genes which may define key driver events in carcinogenesis or potential targets for treatment. We and others have previously shown that in addition to this gene centered or locus centered approach, the structural changes provide important information for classification and survival prediction [8, 9, 26]. The methodology presented in this study complements gene specific analyses by providing a systematic framework to characterize the information embedded in the copy number profile of a tumor. CARMA assigns scores to any predefined collection of genomic regions. In this study, a region was either the whole genome or a chromosome arm. Irrespective of the selection of regions on which to assign scores, the fact that regions are identical across tumors allows CARMA scores to be used directly as features in clustering, regression and classification. Normally, the number of features will also be quite small, thus substantially reducing statistical problems related to high dimensionality.

Molecular taxonomy of breast cancer based on gene expression has proved important for the biological understanding of the disease [16]. IntClust [1] is a more recent driver-based classification of breast cancer and has been shown to also reflect degree of chemosensitivity [27]. The CARMA scores revealed distinct aberration signatures for the 10 IntClust groups, suggesting that the copy number motifs reflect a driver-based classification of tumors. As seen from the Manhattan plots, the expression signatures defining the IntClust subtypes are to a large degree correlated to focal copy number aberrations, representing driver alterations in these subtypes. The copy number aberrations in these driver regions also exhibit differences in their pattern. This is for instance illustrated by the different types of copy number gains found on the 1q arm in the IntClust 8 subtype, as compared to the gains found on the 11q arm in the IntClust2 group. The first type of gain represents non-complex low-amplicon whole arm translocations – captured by the AMP and ASM scores, while the latter represents more complex rearrangements with high-amplicon gains [28] captured by all of the CARMA scores. Even though both of the observed patterns represent copy number gains, the underlying mechanisms causing these patterns are fundamentally different. The CARMA scores manage to capture these nuances, illustrating the potential of the method to discriminate between a richer set of aberrational patterns. The plot also gives an indication of the global background variation from copy number aberrations – maybe most apparent in the IntClust 10 subtype. Interestingly, the degree to which the different subtypes are affected by this background variation seems to correlate well with the fraction of *TP53* mutations observed within each subtype [29]. This again supports the notion that copy number motifs reflect underlying biological traits.

In order to assess the ability of the method to predict breast cancer specific survival, a univariate Cox regression model was fitted to genome-wide CARMA scores in the METABRIC cohort. All genome-wide scores showed a strong and significant association to survival. As a first step this supports the assumption that each of the selected scores are informative and thus qualifies for use in further survival analyses. The scores were combined to produce the unweighted and weighted prognostic indices CPI and CPI_*weighted*_. When these prognostic indices were compared to established clinical and biological parameters through univariate and pairwise bivariate Cox regression analyses, it was evident that the CPI consistently outperformed all other variables. This might point towards a role of specific aberration motifs, proceeding from specific types of genomic instability, as determinants of malignancy potential in a tumor. One shall also take note of the fact that the CPI outperformed GII in these analyses, supporting the idea of additional information added through multifaceted measurements of copy number aberrations.

The observation that the CPI_*weighted*_ obtained poorer prognostic predictions than CPI might stem from the somewhat strict variable selection exerted by the Lasso regression model. The Lasso model excludes arm-specific scores that individually do not contribute strongly to the survival prediction. Aggregated however, these arm-specific scores might confer additional prognostic information. The CPI, which is based on combining all arm-scores in an unweighted manner is not subject to the same kind of selection bias. The fact that this more “inclusive” approach performed better in our analyses suggests that all parts of the genome copy number aberration profile contribute to the real signal when assessing survival. This supports the notion that our method captures omnipresent background variation caused by underlying DNA disruptions.

In the future it would be of high interest to apply the methodology to different cancer types to compare aberration patterns across tumors at different sites, for example using The Cancer Genome Atlas Pan-Cancer dataset [30]. Translocation of genomic material is not captured by any array based DNA analysis, and data from high-throughput sequencing would be required to fully characterize genomic architecture. The complex patterns described in this manuscript are likely to reflect specific mutational processes that could be further elucidated in future studies, linking CARMA with sequencing data. Finally, ASCAT has recently been implemented for whole genome sequencing data [31], and it would be interesting to apply our methodology directly to the allele-specific copy number profiles extracted from such data.

Several extensions of the CARMA algorithm are possible. One could for example increase the genomic resolution by partitioning the genome into a fairly large number of equal-sized regions (say 1000), and then assign separate scores to each of these. At some point, however, the regions may become too small to meaningfully assign scores, most notably for the indices reflecting complex re-arrangements (*STP* and *CRV*). Another possible extension would be to consider regions harboring genes involved in specific processes or pathways, thus directly linking CARMA scores to biological function.

## Author contributions

AVP, GN and OCL performed the statistical and bioinformatical analyses, with contributions from ØB, KL, OMR, MRA, AM, DCW, PVL and HGR. AVP, GN, HGR and OCL developed the CARMA method. VV and AF performed the IntClust subtyping in the Oslo2 cohort. AL and ALBD performed and planned laboratory experiments. OSBREAC and ALBD provided patient materials. AVP, GN, HGR and OCL wrote the manuscript. HGR, OCL, GN and AVP conceived the study. All authors performed critical revision of the manuscript and have read and accepted the final version.

## Acknowledgments

The authors thank all the women who have contributed to this study by donating tumor tissue and blood. We thank Hans Kristian Moen Vollan for vital input and support in the development of the CARMA algorithms. We thank Sandra Jernstrøm for her assistance in preparing the gene expression data on Oslo2, Einar Rødland for his assistance in normalization of the gene expression data on Oslo2, and Phoung Vu, Veronica Skarpeteig, Inger Riise Bergheim and Anja Valen for the *TP* 53 sequencing of Oslo2. David Wedge is supported by the Li Ka Shing Foundation and National Institute for Health Research Oxford Biomedical Research Centre. Peter Van Loo is supported by the Francis Crick Institute, which receives its core funding from Cancer Research UK (FC001202), the UK Medical Research Council (FC001202), and the Wellcome Trust (FC001202). Peter Van Loo is a Winton Group Leader in recognition of the Winton Charitable Foundation’s support towards the establishment of The Francis Crick Institute.

