## Supplementary for "Copy number motifs expose genome instability type and predict driver events and disease outcome in breast cancer"

### Supplementary Material and Methods

### Data material

#### The Oslo2 cohort

A total of 333 patients from the Oslo2 cohort were included in this study. Of these 19 non-invasive were excluded, while 38 samples did not return an ASCAT solution. This left 276 samples for further analyses. The Oslo2 study is an ongoing study where consecutive patients with primary operable (cT1-cT2) breast cancer are included at collaborating hospitals in southeast Norway. Patients were included at the time of surgery after informed consent. All patients in this study were included at Oslo University Hospital.

As described in a previous work [1, 2], tumor material was fresh frozen at - 80° C after macroscopic evaluation of the surgical specimen by an experienced pathologist. The Regional Committee for Medical and Health Research Ethics for southeast Norway have approved the study (approval number 1.2006.1607, amendment 1.2007.1125). Clinical information was collected from hospital records and histopathological data from routine assessment of the tumor. An overview of clinical and pathological variables is given in Supplementary Table 1-2. Fresh frozen tumor was cut with scalpel. One piece of the tumor tissue was used for DNA isolation using the Maxwell® 16 instrument (Promega, USA) and the Maxwell® 16 tissue DNA Purification Kit (Promega).

DNA was isolated according to manufacturer's protocol. In brief, tumor tissue was transferred into the Maxwell cartridge cassettes predispensed with magnetic beads, lysis buffer, and wash buffers of isopropanol and ethanol. The isolation procedure is automated, starting with sample lysis and tissue homogenization, followed by bead isolation of DNA, and finally the washing steps. The DNA was eluted in 200-600 l TE buffer (PH 8.5). DNA is stored at - 20° C. DNA concentration and quality were measured using Nanodrop® ND-1000 (NanoDrop Technologies, USA), which determines the absorbance in the sample by spectrophotometer. 1.5$\mu$l of DNA in solution was used for measurement. The quality of the samples (260/280 absorbance) ranged from 1.33- 2.14 (mean 1.86), the 260/230 absorbances ranged from 0.06-2.39 (mean 1.79). Total RNA isolation was performed using TRIzol (Invitrogen, Life Technologies Corporation, CA, USA) as described previously [3]. Tumor DNA was hybridized to Affymetrix SNP 6.0 arrays per the manufacturer's instructions (Affymetrix, Santa Clara, CA) at AROS Applied Biotechnology (Aarhus, Denmark). Samples that met the quality control criteria established by AROS were subject to further in-house quality assessment.

The mRNA expression was determined by SurePrint G3 Human GE 8·60K one-color microarrays (Agilent, Santa Clara, CA, USA) according to manufacturer's protocol (One-Color Microarray-Based Gene Expression Analysis, Low Input Quick Amp Labeling, v.6.5, May 2010). 100ng of RNA per sample was amplified and hybridized on array. The array includes 42,405 unique 60-mer probes, targeting 27,958 Entrez Gene RNAs and 7,419 lincRNAs. Scanning was performed using Agilent Scanner G2565A and AgilentG3_GX_1Color was used as profile. The signals were extracted using FeatureExtraction v.10.7.3.1 and protocol GE1_107_Sep09. The data was quantile normalized, hospital-centered and log2-transformed [1].

#### The METABRIC cohort

The Molecular Taxonomy of Breast Cancer International Consortium (METABRIC) cohort consists of 1980 patients with primary breast cancer [4]. Fresh frozen tumor tissue were collected from tumor banks in the UK (Nottingham, Addenbrooke's in Cambridge and Guys hospital in London) and Canada (Vancouver and Manitoba). Copy number (Affymetrix SNP 6.0) and gene expression (Illumina HT-12 v3 Expression Beadchip) were available and deposited to the European Genome-Phenome Archive (EGA, http://www.ebi.ac.uk/ega/) hosted by the European Bioinformatics Institute (EBI, Hinxton, UK) with accession number EGAS00000000083. The expression data were processed as described by [4] and the raw intensities were quantile normalized and log2-transformed.

A total of 30 samples were excluded due to unknown, pre-invasive or benign histology. Seven samples did not return an ASCAT solution, leaving 1943 samples for further analyses. A random partition of the dataset (2:1 randomization) into a discovery set (n=1295) and a test set (n=648) was performed.

Extensive clinical annotation is available for these patients. Median follow-up time for the full set is 7.3 years. Tumor size were categorized according to AJCC guidelines [5]. Estrogen receptor status (ER) by immunohistochemistry (IHC) was available for 1914 cases (98.5%), the 29 other samples were scored by the expression value of *ESR1* as described in [6]. HER2 status status by IHC was not available and hence scored by the expression value of *HER2* [6]. The clinical variables are presented in Supplementary Table 1-2.

#### The OsloVal cohort

The OsloVal cohort is a historic archive material from the Norwegian Radium Hospital, Oslo, Norway, where excess tumor material was stored after biochemical ER assay. For a detailed description of this data cohort, confer [7]. The Regional Committee for Medical and Health Research Ethics for southeast Norway have approved the study (approval number 2010/498). DNA and RNA were isolated from 184 samples and profiled on Affymetrix SNP 6.0 arrays and Illumina HT12 Bead Chip. Three samples were excluded due to unknown/uncertain histology and 16 samples did not have an ASCAT solution, leaving 165 samples for further analysis. Clinical and pathological variables are given in Supplementary Table 1-2. The original ER-status obtained for the OsloVal samples was not used, as it involved different methods being used on different samples. Instead, the ER-status was determined by considering the expression of ESR1. Examination of the distribution of ESR1 expression values across the cohort revealed two distinct peaks separated by a trough at $ESR1 \approx 7.5$, and based on this we called tumors with $ESR1 < 7.5$ as ER-negative and tumors with $ESR1 \geq 7.5$ as ER-positive. The copy number profiles of the samples in the OsloVal cohort were more affected by noise than the two other datasets, with numerous very short, scattered low-level copy number changes across the whole genome. As a consequence, we filtered out segments that were shorter than 5Mb and differed by no more than two copies from its neighboring segments. This approach proved to perform well for removing short spikes, while preserving more complex rearrangements like firestorms.

#### Regional instability score algorithm

We now describe the algorithm for computation of the six regional instability scores defined in the previous section. The region can be chosen as any contiguous section of a chromosome. In this paper, we focus on the case where regions correspond to chromosome arms. With 42 chromosome arms(excluding the short arms of the acrocentric chromosomes) we thus obtain $6 \times 42 = 252$ score values per tumor sample.

Suppose the genomic region has length $L$, and let the associated allele-specific copy number profile be $(n_{Ai}, n_{Bi})$, $i = 1, \ldots, m$. Let $p_i$ be the physical location of the $i$th probe and $s_i = p_i/L$ the corresponding mapped location of the $i$th probe in $R = [0, 1]$. Whenever $n_{Ak} + n_{Bk} \neq n_{A,k+1} + n_{B,k+1}$, we define $(s_k + s_{k+1})/2$ to be a change point. Let $0 < t_1 < t_2 < \cdots < t_{r-1} < 1$ be all the change points, and define $t_{-2} = t_{-1} = t_0 = 0$ and $t_r = t_{r+1} = t_{r+2} = 1$. The total copy number is then

$$f(t) = n_{Ak_i} + n_{Bk_i}, \quad t \in [t_{i-1}, t_i) \tag{1}$$

for $i = 1, \ldots, r$, where $k_i$ satisfies $s_{k_i} \in [t_{i-1}, t_i)$. In the following, the function $f(t)$ will be represented by the parameterization $f_0 = 0$, $f_i = f(t_{i-1})$ $(i = 1, \ldots, r)$, $f_{r+1} = 0$. Now recall that $Df(t)$ is the slope of the line segment connecting the pair of segment centers immediately to the left and right of position $t$. Defining $t_i^{(1)} = (t_{i-1} + t_i)/2$ for $i = -1, 0, \ldots, r + 2$, we thus have

$$Df(t) = \frac{f_i - f_{i-1}}{t_i^{(1)} - t_{i-1}^{(1)}}, \quad t \in [t_{i-1}^{(1)}, t_i^{(1)}) \tag{2}$$

for $i = 1, \ldots, r + 1$. In the following, the function $Df(t)$ will be represented by the parameterization $(Df)_0 = 0$, $(Df)_i = Df(t_{i-1}^{(1)})$ $(i = 1, \ldots, r + 1)$, $(Df)_{r+2} = 0$. We define $D^2 f(t)$ in a similar manner by first letting $t_i^{(2)} = (t_{i-1}^{(1)} + t_i^{(1)})/2$ for $i = 0, \ldots, r + 1$, and

$$D^2 f(t) = \frac{(Df)_i - (Df)_{i-1}}{t_i^{(2)} - t_{i-1}^{(2)}}, \quad t \in [t_{i-1}^{(2)}, t_i^{(2)}) \tag{3}$$

for $i = 1, \ldots, r + 2$. In the following, the function $D^2 f(t)$ will be represented by the parameterization $(D^2 f)_i = D^2 f(t_{i-1}^{(2)})$ $(i = 1, \ldots, r + 1)$, $(D^2 f)_{r+2} = 0$.

Evaluation of the scores $J_1, \ldots, J_6$ involves integration of piecewise constant functions, and can thus be expressed as finite sums over the segments.

We have

$$J_1 = \sum_{i=1}^{r} (t_i - t_{i-1})(f_i)_+^2 \tag{4}$$

$$J_2 = \sum_{i=1}^{r} (t_i - t_{i-1})(f_i)_-^2 \tag{5}$$

$$J_3 = \sum_{i=1}^{r+1} (t_i^{(1)} - t_{i-1}^{(1)})(Df)_i^2 = \sum_{i=1}^{r+1} \frac{(f_i - f_{i-1})^2}{t_i^{(1)} - t_{i-1}^{(1)}} \tag{6}$$

$$J_4 = \sum_{i=1}^{r+2} (t_i^{(2)} - t_{i-1}^{(2)})(D^2 f)_i^2 = \sum_{i=1}^{r+2} \frac{((Df)_i - (Df)_{i-1})^2}{t_i^{(2)} - t_{i-1}^{(2)}} \tag{7}$$

To insure robustness against outliers, $\psi_R^{(1)}$ and $\psi$ were calculated as the median rather than the mean of the regional and genome-wide copy number distribution. In most cases, this will have a small effect, however, as it is our experience that the median is usually close to the mean for the type of data considered here. The scores in equations (6) and (7) are weighted by the size of the genomic region. These scores capture focal complex events of a size that do not necessarily reflect the size of the genomic region, hence weighting is required to counteract the normalization of the genomic scale to the interval $[0, 1]$. In addition, to ensure that the values of these two indices are not unduly influenced by very small distances between break points, each term in the right hand sides of these equations is dampened by application of a soft threshold function $x \mapsto \tanh(x)$. Specifically, we have $J_2 = \sum (f_i - f_{i-1})^2 \cdot \tanh(1/(t_i^{(1)} - t_{i-1}^{(1)}))$ and $J_3 = \sum ((Df)_i - (Df)_{i-1})^2 \cdot \tanh(1/(t_i^{(2)} - t_{i-1}^{(2)}))$. Evaluation of $J_6$ requires allele-specific copy number values. Whenever $n_{Ak} \neq n_{A,k+1}$ or $n_{Bk} \neq n_{B,k+1}$, we define $(s_k + s_{k+1})/2$ to be a change point. Denoting change points $0 < t_1 < t_2 < \cdots < t_{r-1} < 1$, define $t_0 = 0$ and $t_r = 1$ and observe that the number of A-alleles and B-alleles is

$$f_A(t) = n_{Ak_i} \text{ and } f_B(t) = n_{Bk_i} \quad t \in [t_{i-1}, t_i) \tag{8}$$

for $i = 1, \ldots, r$, where $k_i$ satisfies $s_{k_i} \in [t_{i-1}, t_i)$. Let $f_A(t)$ be represented by the parameterization $f_{A0} = 0$, $f_{Ai} = f_A(t_{i-1})$ $(i = 1, \ldots, r)$, $f_{A,r+1} = 0$, and let $f_B(t)$ be represented by $f_{B0} = 0$, $f_{Bi} = f_B(t_{i-1})$ $(i = 1, \ldots, r)$, $f_{B,r+1} = 0$. Then,

$$J_5 = \sum_{i=1}^{r} (t_i - t_{i-1}) 1_0(f_{Bi}) \tag{9}$$

$$J_6 = \sum_{i: f_{Ai}=0} (t_i - t_{i-1})(f_{Ai} - f_{Bi})^2 \tag{10}$$

#### Standardization of scores

All six scores were $\log_2$-transformed and normalized by dividing by the 99th percentile. To have a common reference, the 99th percentile in the METABRIC discovery set were used to standardize all data sets.

#### Survival analyses

In the univariate analyses we applied Cox proportional hazard regression using the function `coxph` in the `survival` package. For the multivariate analyses, we applied Cox regression with

Lasso penalty [8] as implemented in the `glmnet` package [9, 10]. The lasso is a regularization method which shrinks regression coefficients towards zero by putting an upper bound on the $L_1$-norm of the coefficients (i.e. $\sum_{j=1}^{p} |\beta_j| \leq \lambda$) in the maximization of the partial log likelihood. The amount of shrinkage is determined by the tuning parameter $\lambda$, which can be estimated by cross-validation. Typically, one selects the value of $\lambda$ that gives the minimum mean cross-validation error, however, we observed that the selected value depended on the fold assignment. We ran n-fold cross-validation(leave one out) 100 times and each time selected the value of $\lambda$ that gave the minimum mean cross-validation error; we then chose the 95'th percentile among this set of $\lambda$ values to use in the final lasso model. The weighted prognostic index ($\text{CPI}_{weighted}$) was calculated as $\text{CPI}_{weighted} = \boldsymbol{x}_i^T \hat{\boldsymbol{\beta}}$, where $\boldsymbol{x}_i$ represents the CARMA arm-scores for patient $i$ in the validation data set and $\hat{\boldsymbol{\beta}}$ are the estimated coefficients in the survival prediction models found for the discovery set.

#### GII

The Genomic Instability Index (GII) is the fraction of the genome with aberrant copy number, here defined as an aberration from the median total copy number across the whole genome (denoted $\psi$). Formally, this can be expressed as $GII = \int_R I(|f(t) - \psi| > 0)\, dt$ where $f(t)$ denotes total copy number at locus $t \in R$, $I(.)$ takes values 0 and 1 depending on whether its argument is false or true, and $R = [0, 1]$ denotes positions on the genome.

#### Breast cancer subtyping

PAM50 subtypes were found using the method described in [11] and with the subtype centroids provided on the accompanying web site. In brief, for each sample, the gene expression values for the 50 genes in the PAM50 gene list were extracted. Three genes in the PAM50 signature were replaced with synonyms in our data, namely *CDCA1* with *NUF2*, *KNTC2* with *NDC80*, and *ORC6L* with *ORC6*. When multiple probes with identical PAM50 gene identifier was found, the average probe value was used for Agilent expression data and what was considered to be the probe least affected by variant splicing was used for Illumina expression data. It is recommended in [11] to perform an initial test-to-train set normalization, and for this purpose two centroids were calculated by averaging gene expression values over all ER-positive samples and all ER-negative samples, respectively. Clinical ER status obtained through immunohistochemical staining of ER was used. A combined centroid was next defined as a weighted average of the ER-negative centroid and the ER-positive centroid, the weights being c and 1-c, where c is the proportion of ER-negative samples in the original data set (the training data set) used to define the PAM50 centroids. The samples to be subtyped were then centered by aligning the combined centroid with the centroid of the training data set, achieved by subtracting the combined centroid from the expression vector of each sample and then adding the centroid of the training data set. Finally, a subtype label (Luminal A, Luminal B, Her2-enriched, Basal-like, or Normal-like) was assigned to each sample by calculating the Spearman correlation between the sample's expression vector and each of the five PAM50 centroids and selecting the one with the highest correlation.

IntClust subtypes were determined by the approach described in [4], which is based on 754 features (39 segmented copy number features and 715 gene expression values). Features were first matched, using genomic position or gene name for copy number features and gene name for expression features. To make features comparable to those in the original dataset, they were normalized as outlined in [4]. Finally, samples were assigned to the 10 classes by the Nearest Shrunken Centroids method [12], using the original centroids and within-cluster standard deviations of each of the 754 features

Supplementary Table 1 provides an overview of the distribution of the PAM50 subtypes and the IntClust subtypes in each dataset.

### Supplementary Figures

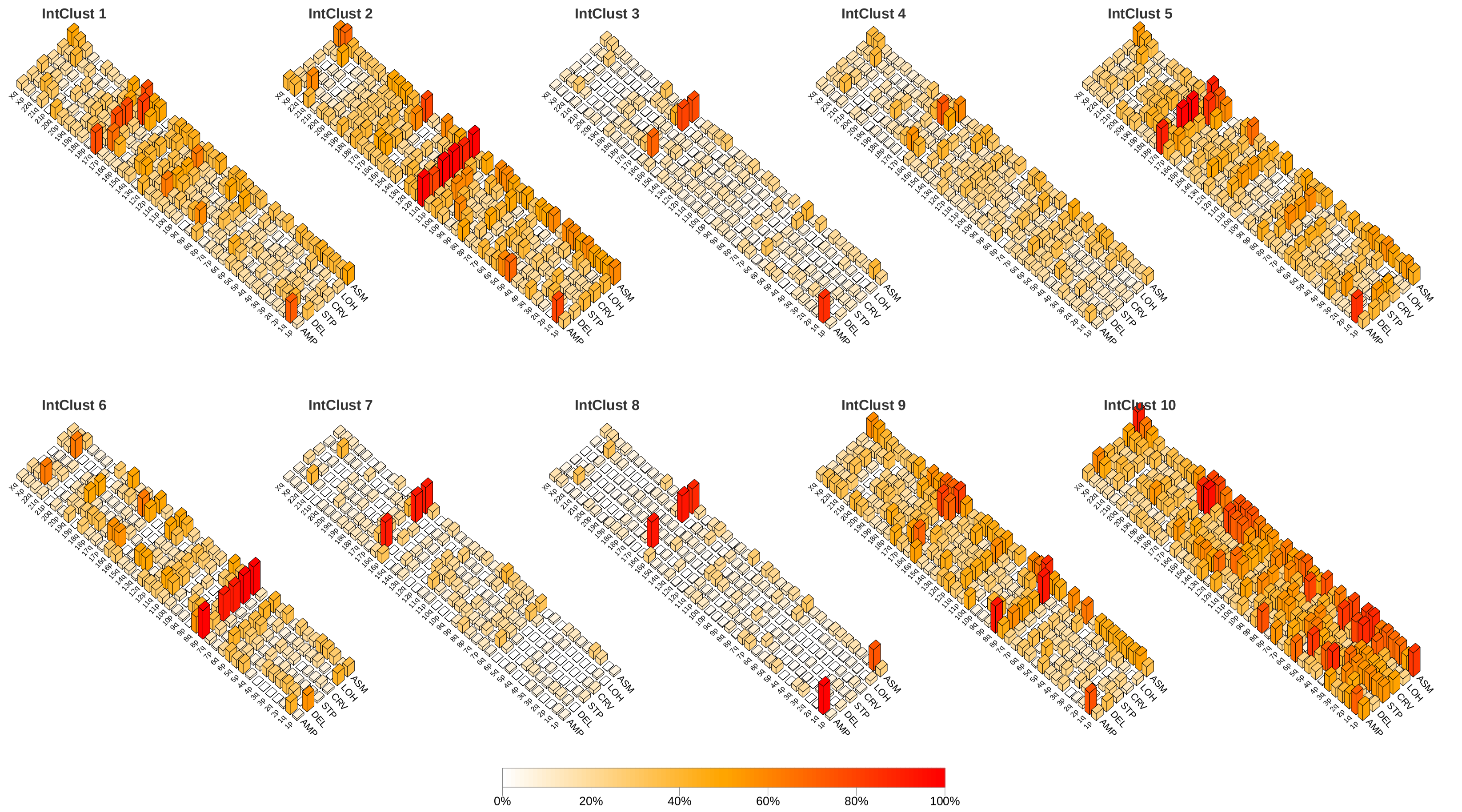


**Figure S1.** Score landscapes for the IntClust subtypes in the Oslo2 cohort. The bars reflect, for each of the 6 CARMA indices and each chromosome arm, the percentage of tumors in the Oslo2 cohort that have a score larger than the index median (calculated across all arms and ignoring zero scores within each of the 6 CARMA indices). See Supplementary Table 1 for the number of tumors in each IntClust subtype.

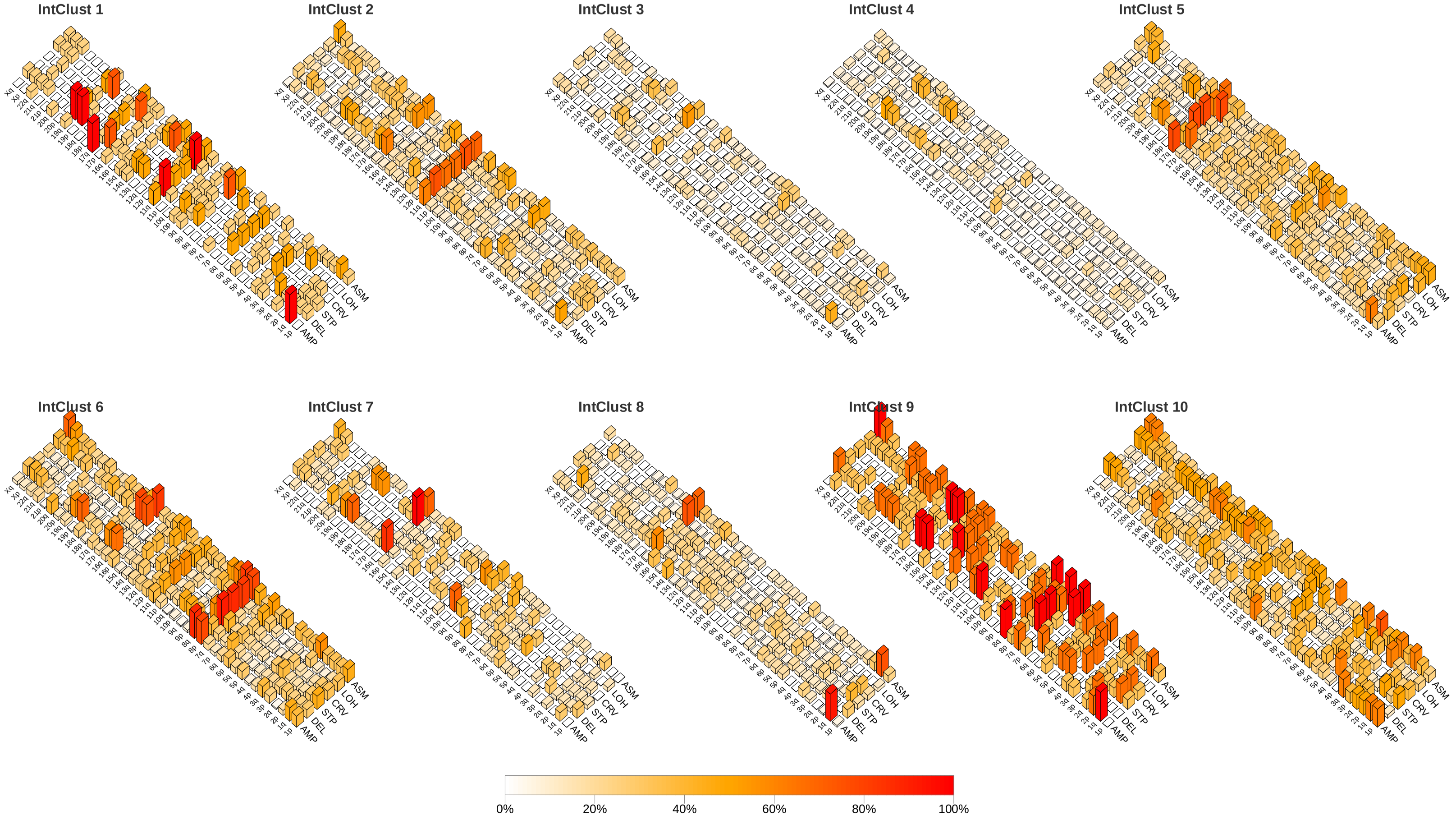


**Figure S2.** Score landscapes for the IntClust subtypes in the OsloVal cohort. The bars reflect, for each of the 6 CARMA indices and each chromosome arm, the percentage of tumors in the OsloVal cohort that have a score larger than the index median (calculated across all arms and ignoring zero scores within each of the 6 CARMA indices). See Supplementary Table 1 for the number of tumors in each IntClust subtype.

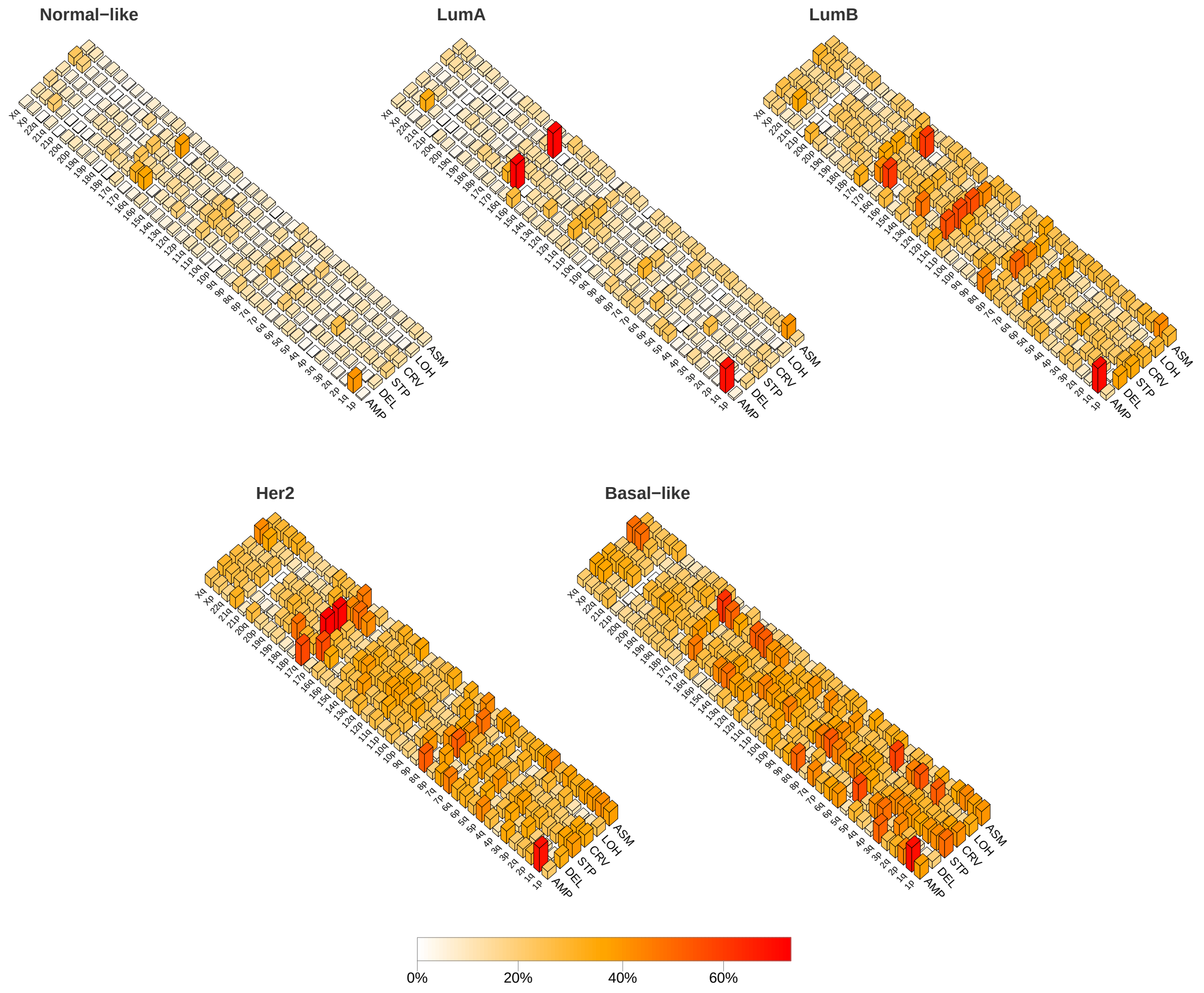


**Figure S3.** Score landscapes for PAM50 subtypes in the METABRIC cohort. The bars reflect, for each of the 6 CARMA indices and each chromosome arm, the percentage of tumors in the METABRIC set that have a score larger than the index median (calculated across all arms and ignoring zero scores within each of the 6 CARMA indices).

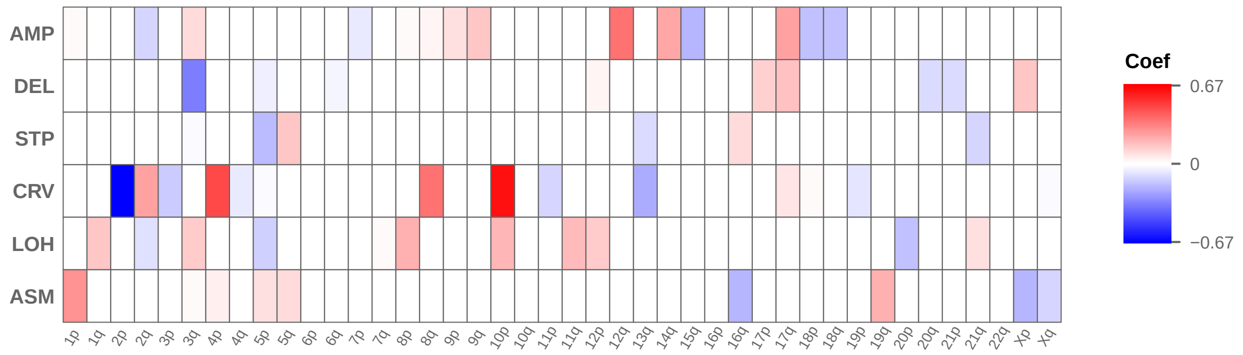


**Figure S4.** Heatmap giving arm-specific coefficients from Lasso-Cox analysis of disease specific survival in the METABRIC discovery set.

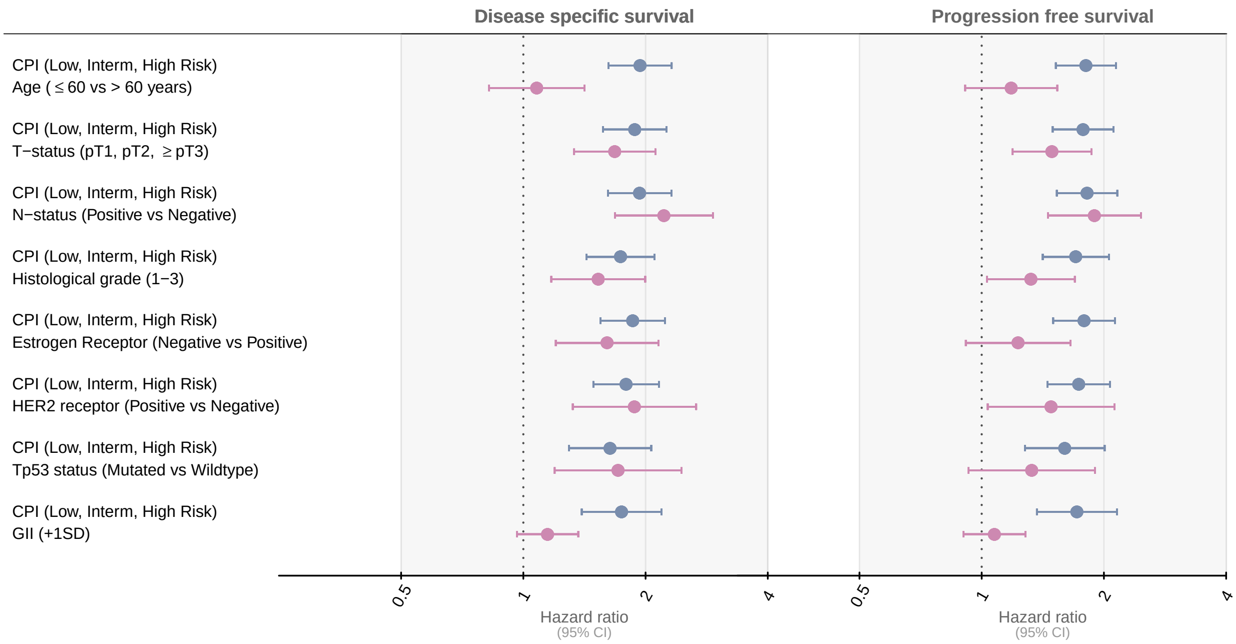


**Figure S5.** Hazard ratios and 95% confidence intervals(CI) for clinical variables, the CARMA Prognostic Index(CPI) based on unweighted averages of arm-wise CARMA scores, and the genomic instability index(GII) in the METABRIC test set. Shown are adjusted estimates for disease specific survival (DSS) and progression free survival (PFS).

### Supplementary Tables

**Table S1.** Subtype distribution of the four study cohorts (METABRIC discovery, METABRIC test, Oslo2 and OsloVal). Shown is the number and proportion of samples in each cohort that were assigned to each of the five PAM50 subtypes and each of the ten IntClust subtypes.

| | METABRIC discovery | METABRIC test | Oslo2 | OsloVal |
|---|---|---|---|---|
| No of samples | 1295 | 648 | 276 | 147 |
| PAM50 subtypes | | | | |
| Luminal A | 461 (35.6) | 249 (38.4) | 107 (38.8) | 29 (19.7) |
| Luminal B | 323 (24.9) | 158 (24.4) | 63 (22.8) | 47 (32) |
| Basal-like | 215 (16.6) | 106 (16.4) | 36 (13) | 26 (17.7) |
| Her2-enriched | 159 (12.3) | 77 (11.9) | 31 (11.2) | 21 (14.3) |
| Normal-like | 133 (10.3) | 56 (8.6) | 15 (5.4) | 24 (16.3) |
| IntClust subtypes | | | | |
| IntClust1 | 91 (7) | 47 (7.3) | 22 (8) | 4 (2.7) |
| IntClust2 | 41 (3.2) | 30 (4.6) | 10 (3.6) | 21 (14.3) |
| IntClust3 | 190 (14.7) | 97 (15) | 45 (16.3) | 11 (7.5) |
| IntClust4 | 216 (16.7) | 113 (17.4) | 40 (14.5) | 26 (17.7) |
| IntClust5 | 124 (9.6) | 60 (9.3) | 22 (8) | 19 (12.9) |
| IntClust6 | 62 (4.8) | 22 (3.4) | 14 (5.1) | 24 (16.3) |
| IntClust7 | 122 (9.4) | 65 (10) | 15 (5.4) | 7 (4.8) |
| IntClust8 | 195 (15.1) | 101 (15.6) | 32 (11.6) | 17 (11.6) |
| IntClust9 | 95 (7.3) | 48 (7.4) | 17 (6.2) | 3 (2) |
| IntClust10 | 159 (12.3) | 65 (10) | 25 (9.1) | 8 (5.4) |

**Table S2.** Overview of clinical and pathological variables. Survival data were not available for the Oslo2 cohort; HER2 status and Tp53 status were not available for the OsloVal cohort.

| | METABRIC discovery | METABRIC test | Oslo2 | OsloVal |
|---|---|---|---|---|
| No. of samples | 1295 | 648 | 276 | 147 |
| Age at diagnosis (median) | 62 | 61.2 | 56.8 | 59 |
| T-status | | | | |
| pT1 | 557 (43) | 285 (44) | 150 (54.3) | 57 (38.8) |
| pT2 | 664 (51.3) | 321 (49.5) | 110 (39.9) | 58 (39.5) |
| pT3 | 62 (4.8) | 34 (5.2) | 12 (4.3) | 9 (6.1) |
| pT4 | 0 (0) | 0 (0) | 0 (0) | 16 (10.9) |
| N-status | | | | |
| pN- | 674 (52) | 343 (52.9) | 172 (62.3) | 81 (55.1) |
| pN+ | 621 (48) | 305 (47.1) | 104 (37.7) | 58 (39.5) |
| Histological grade | | | | |
| Grade 1 | 115 (8.9) | 50 (7.7) | 40 (14.5) | 11 (7.5) |
| Grade 2 | 519 (40.1) | 244 (37.7) | 116 (42) | 59 (40.1) |
| Grade 3 | 614 (47.4) | 331 (51.1) | 119 (43.1) | 41 (27.9) |
| Estrogen receptor | | | | |
| Positive | 993 (76.7) | 487 (75.2) | 222 (80.4) | 114 (77.6) |
| Negative | 282 (21.8) | 152 (23.5) | 54 (19.6) | 33 (22.4)) |
| Progesterone receptor | | | | |
| Positive | 680 (52.5) | 344 (53.1) | 196 (71) | 72 (49) |
| Negative | 615 (47.5) | 304 (46.9) | 80 (29) | 63 (42.9) |
| HER2 status | | | | |
| Positive | 164 (12.7) | 76 (11.7) | 26 (9.4) | – |
| Negative | 1131 (87.3) | 572 (88.3) | 250 (90.6) | – |
| Tp53 status | | | | |
| Wildtype | 692 (53.4) | 332 (51.2) | 183 (66.3) | – |
| Mutation | 265 (20.5) | 125 (19.3) | 91 (33) | – |
| Survival analyses events | | | | |
| BC deaths | 428 (33.1) | 212 (32.7) | – | 58 (39.5) |
| All deaths | 752 (58.1) | 375 (57.9) | – | 104 (70.7) |

**Table S3.** Univariate Cox regression on the six whole-genome CARMA scores found by unweighted averaging of the arm-wise scores. Results are shown for disease specific survival (DSS) in the METABRIC cohort (n = 1943). Shown is the estimated hazard ratio, 95 % confidence intervals for the hazard ratio, Z-score and P-value. Results are also shown for a model using the Genomic Instability Index (GII) as predictor.

| Score | HR | Lower 95% CI | Upper 95% CI | Z-score | P-value |
|---|---|---|---|---|---|
| AMP | 5.55 | 3.52 | 8.76 | 7.37 | 2e-13 |
| DEL | 2.77 | 1.87 | 4.10 | 5.10 | 3e-07 |
| STP | 4.15 | 3.05 | 5.66 | 9.01 | <2e-16 |
| CRV | 3.35 | 2.58 | 4.37 | 8.97 | <2e-16 |
| LOH | 3.56 | 2.26 | 5.60 | 5.49 | 4e-08 |
| ASM | 8.05 | 4.55 | 14.22 | 7.18 | 1e-12 |
| GII | 2.95 | 2.24 | 3.89 | 7.66 | 1e-14 |

**Table S4.** Prognostic value of clinical variables, the CPI and $CPI_{weighted}$, and the genomic instability index (GII) for disease specific survival (DSS) in the METABRIC test set (n = 648). HR: hazard ratio.

| Type | Covariate | P value | HR | Lower 95% CI | Upper 95% CI |
|---|---|---|---|---|---|
| Unadjusted | Age ($\leq$ 60 vs > 60 years) | 2.2e-01 | 1.19 | 0.91 | 1.55 |
| Unadjusted | T-status (pT1, pT2, $\geq$pT3) | 9.2e-07 | 1.76 | 1.40 | 2.20 |
| Unadjusted | N-status (Positive vs Negative) | 5.5e-09 | 2.28 | 1.73 | 3.01 |
| Unadjusted | Histological grade (1-3) | 5.9e-08 | 2.01 | 1.56 | 2.58 |
| Unadjusted | Estrogen Receptor (Negative vs Positive) | 2.0e-06 | 2.00 | 1.50 | 2.66 |
| Unadjusted | HER2 receptor (Positive vs Negative) | 1.6e-09 | 2.79 | 2.00 | 3.89 |
| Unadjusted | Tp53 status (Mutated vs Wildtype) | 7.2e-08 | 2.44 | 1.77 | 3.38 |
| Unadjusted | GII ($+1SD$) | 1.2e-09 | 1.51 | 1.32 | 1.73 |
| Unadjusted | CPI (Low,Interm.,High risk) | 2.4e-13 | 1.95 | 1.63 | 2.33 |
| Unadjusted | CPI weighted (Low,Interm.,High risk) | 2.2e-07 | 1.57 | 1.32 | 1.87 |
| Adjusted | CPI (Low,Interm.,High risk) | 4.5e-13 | 1.94 | 1.62 | 2.32 |
| | Age ($\leq$ 60 vs > 60 years) | 5.8e-01 | 1.08 | 0.82 | 1.42 |
| Adjusted | CPI (Low,Interm.,High risk) | 6.8e-12 | 1.88 | 1.57 | 2.25 |
| | T-status (pT1, pT2, $\geq$pT3) | 1.0e-05 | 1.68 | 1.33 | 2.11 |
| Adjusted | CPI (Low,Interm.,High risk) | 7.3e-13 | 1.93 | 1.62 | 2.32 |
| | N-status (Positive vs Negative) | 1.8e-08 | 2.22 | 1.68 | 2.93 |
| Adjusted | CPI (Low,Interm.,High risk) | 2.0e-08 | 1.74 | 1.43 | 2.10 |
| | Histological grade (1-3) | 1.8e-03 | 1.53 | 1.17 | 2.00 |
| Adjusted | CPI (Low,Interm.,High risk) | 2.9e-11 | 1.86 | 1.55 | 2.23 |
| | Estrogen Receptor (Negative vs Positive) | 1.4e-03 | 1.61 | 1.20 | 2.15 |
| Adjusted | CPI (Low,Interm.,High risk) | 8.1e-10 | 1.79 | 1.49 | 2.16 |
| | HER2 receptor (Positive vs Negative) | 4.1e-04 | 1.88 | 1.32 | 2.66 |
| Adjusted | CPI (Low,Interm.,High risk) | 3.7e-05 | 1.64 | 1.29 | 2.07 |
| | Tp53 status (Mutated vs Wildtype) | 3.4e-03 | 1.71 | 1.19 | 2.45 |
| Adjusted | CPI (Low,Interm.,High risk) | 1.4e-06 | 1.75 | 1.39 | 2.19 |
| | GII ($+1SD$) | 1.2e-01 | 1.15 | 0.96 | 1.37 |
| Adjusted | CPI weighted (Low,Interm.,High risk) | 3.4e-07 | 1.56 | 1.32 | 1.86 |
| | Age ($\leq$ 60 vs > 60 years) | 4.3e-01 | 1.12 | 0.85 | 1.46 |
| Adjusted | CPI weighted (Low,Interm.,High risk) | 1.9e-06 | 1.52 | 1.28 | 1.80 |
| | T-status (pT1, pT2, $\geq$pT3) | 4.6e-06 | 1.70 | 1.36 | 2.14 |
| Adjusted | CPI weighted (Low,Interm.,High risk) | 1.4e-06 | 1.53 | 1.29 | 1.81 |
| | N-status (Positive vs Negative) | 3.4e-08 | 2.19 | 1.66 | 2.88 |
| Adjusted | CPI weighted (Low,Interm.,High risk) | 9.1e-04 | 1.36 | 1.13 | 1.63 |
| | Histological grade (1-3) | 3.3e-05 | 1.75 | 1.34 | 2.28 |
| Adjusted | CPI weighted (Low,Interm.,High risk) | 9.6e-06 | 1.49 | 1.25 | 1.77 |
| | Estrogen Receptor (Negative vs Positive) | 1.9e-04 | 1.74 | 1.30 | 2.33 |
| Adjusted | CPI weighted (Low,Interm.,High risk) | 2.5e-05 | 1.46 | 1.22 | 1.74 |
| | HER2 receptor (Positive vs Negative) | 1.5e-06 | 2.32 | 1.65 | 3.27 |
| Adjusted | CPI weighted (Low,Interm.,High risk) | 2.5e-03 | 1.39 | 1.12 | 1.72 |
| | Tp53 status (Mutated vs Wildtype) | 3.1e-05 | 2.06 | 1.47 | 2.90 |
| Adjusted | CPI weighted (Low,Interm.,High risk) | 1.1e-02 | 1.29 | 1.06 | 1.57 |
| | GII ($+1SD$) | 6.9e-05 | 1.37 | 1.17 | 1.61 |

**Table S5.** Prognostic value of clinical variables, the CPI and $CPI_{weighted}$, and the genomic instability index (GII) for progression free survival (PFS) in the METABRIC test set (n = 648). HR: hazard ratio.

| Type | Covariate | P value | HR | Lower 95% CI | Upper 95% CI |
|---|---|---|---|---|---|
| Unadjusted | Age ($\leq$ 60 vs > 60 years) | 5.3e-02 | 1.29 | 1.0 | 1.68 |
| Unadjusted | T-status (pT1, pT2, $\geq$pT3) | 2.9e-05 | 1.59 | 1.28 | 1.98 |
| Unadjusted | N-status (Positive vs Negative) | 8.2e-07 | 1.94 | 1.49 | 2.52 |
| Unadjusted | Histological grade (1-3) | 3.2e-06 | 1.74 | 1.38 | 2.20 |
| Unadjusted | Estrogen Receptor (Negative vs Positive) | 2.3e-03 | 1.57 | 1.2 | 2.09 |
| Unadjusted | HER2 receptor (Positive vs Negative) | 7.0e-06 | 2.19 | 1.56 | 3.09 |
| Unadjusted | Tp53 status (Mutated vs Wildtype) | 1.0e-04 | 1.90 | 1.4 | 2.62 |
| Unadjusted | GII (+1$SD$) | 3.9e-08 | 1.43 | 1.26 | 1.63 |
| Unadjusted | CPI (Low,Interm.,High risk) | 3.8e-12 | 1.83 | 1.54 | 2.16 |
| Unadjusted | CPI weighted (Low,Interm.,High risk) | 6.6e-09 | 1.64 | 1.39 | 1.93 |
| Adjusted | CPI (Low,Interm.,High risk) | 1.1e-11 | 1.81 | 1.52 | 2.14 |
| | Age ($\leq$ 60 vs > 60 years) | 2.1e-01 | 1.18 | 0.91 | 1.54 |
| Adjusted | CPI (Low,Interm.,High risk) | 6.2e-11 | 1.78 | 1.50 | 2.11 |
| | T-status (pT1, pT2, $\geq$pT3) | 4.7e-04 | 1.49 | 1.19 | 1.86 |
| Adjusted | CPI (Low,Interm.,High risk) | 8.7e-12 | 1.82 | 1.53 | 2.16 |
| | N-status (Positive vs Negative) | 2.0e-06 | 1.90 | 1.46 | 2.47 |
| Adjusted | CPI (Low,Interm.,High risk) | 2.5e-08 | 1.71 | 1.41 | 2.06 |
| | Histological grade (1-3) | 2.8e-02 | 1.32 | 1.03 | 1.70 |
| Adjusted | CPI (Low,Interm.,High risk) | 8.7e-11 | 1.79 | 1.50 | 2.13 |
| | Estrogen Receptor (Negative vs Positive) | 1.7e-01 | 1.23 | 0.91 | 1.65 |
| Adjusted | CPI (Low,Interm.,High risk) | 1.2e-09 | 1.73 | 1.45 | 2.07 |
| | HER2 receptor (Positive vs Negative) | 3.2e-02 | 1.48 | 1.03 | 2.12 |
| Adjusted | CPI (Low,Interm.,High risk) | 4.2e-05 | 1.60 | 1.28 | 2.01 |
| | Tp53 status (Mutated vs Wildtype) | 1.2e-01 | 1.33 | 0.93 | 1.90 |
| Adjusted | CPI (Low,Interm.,High risk) | 3.2e-06 | 1.72 | 1.37 | 2.15 |
| | GII (+1$SD$) | 4.2e-01 | 1.07 | 0.90 | 1.28 |
| Adjusted | CPI weighted (Low,Interm.,High risk) | 1.6e-08 | 1.62 | 1.37 | 1.91 |
| | Age ($\leq$ 60 vs > 60 years) | 1.6e-01 | 1.21 | 0.93 | 1.57 |
| Adjusted | CPI weighted (Low,Interm.,High risk) | 4.6e-08 | 1.60 | 1.35 | 1.90 |
| | T-status (pT1, pT2, $\geq$pT3) | 2.6e-04 | 1.51 | 1.21 | 1.88 |
| Adjusted | CPI weighted (Low,Interm.,High risk) | 1.8e-08 | 1.61 | 1.37 | 1.90 |
| | N-status (Positive vs Negative) | 2.3e-06 | 1.89 | 1.45 | 2.46 |
| Adjusted | CPI weighted (Low,Interm.,High risk) | 2.3e-05 | 1.47 | 1.23 | 1.76 |
| | Histological grade (1-3) | 2.1e-03 | 1.47 | 1.15 | 1.87 |
| Adjusted | CPI weighted (Low,Interm.,High risk) | 5.9e-08 | 1.61 | 1.36 | 1.91 |
| | Estrogen Receptor (Negative vs Positive) | 9.9e-02 | 1.28 | 0.95 | 1.73 |
| Adjusted | CPI weighted (Low,Interm.,High risk) | 7.6e-07 | 1.54 | 1.30 | 1.83 |
| | HER2 receptor (Positive vs Negative) | 5.1e-03 | 1.67 | 1.17 | 2.38 |
| Adjusted | CPI weighted (Low,Interm.,High risk) | 1.5e-04 | 1.51 | 1.22 | 1.87 |
| | Tp53 status (Mutated vs Wildtype) | 2.6e-02 | 1.48 | 1.05 | 2.08 |
| Adjusted | CPI weighted (Low,Interm.,High risk) | 7.7e-04 | 1.41 | 1.16 | 1.73 |
| | GII (+1$SD$) | 1.1e-02 | 1.23 | 1.05 | 1.44 |
